# Source Reconstruction Without an MRI using Optically Pumped Magnetometer based Magnetoencephalography

**DOI:** 10.1101/2025.01.28.635329

**Authors:** Natalie Rhodes, Lukas Rier, Elena Boto, Ryan Hill, Matthew J. Brookes

## Abstract

Source modelling in magnetoencephalography (MEG) maps the spatial origins of electrophysiological signals in the brain. Typically, this requires an anatomical MRI scan of the subject’s head, from which a model of the neuromagnetic field (the forward model) is derived. Wearable MEG – based on optically pumped magnetometers (OPMs) – enables MEG measurement from participants who struggle to cope with conventional scanning environments (e.g. children), enabling study of novel cohorts. However, its value is limited if an MRI scan is still required for source modelling. Here we describe a method of warping template MRIs to 3D structured-light scans of the head, to generate “pseudo-MRIs”. We apply our method using data from 20 participants during a sensory task, measuring induced (beta band) responses and whole-brain functional connectivity. Results show that the group average locations of peak task-induced beta modulation were separated by 2.75 mm, when comparing real- and pseudo-MRI approaches. Group averaged time-frequency spectra were also highly correlated (Pearson correlation 0.99) as were functional connectome matrices (0.87), and global connectivity (0.98). In sum, our results demonstrate that source-localized OPM-MEG data, modelled with and without an individual MRI scan can be similar. This will be useful for future studies where MRI data capture is challenging.

## Introduction

Source reconstruction of magnetoencephalography (MEG) data is used to generate 3D images showing the spatial origins of electrophysiological signals across the brain. Usually, this requires an anatomical MRI scan of the subject’s head, alongside accurate knowledge of MEG sensor locations and orientations relative to brain anatomy (from the MRI). These data are combined to generate a volume conductor model, from which a mathematical description of the magnetic fields generated by the brain (known as the *forward model*) is derived. This forward model is then combined with the MEG data to produce source images, and the MRI scan also allows those functional images to be overlaid onto anatomical structure. This technique is commonplace, however, in many studies the acquisition of an MRI scan can be challenging due to the unnatural, noisy and claustrophobic scanning environment, which is not always well-tolerated, particularly by children. Other important considerations include the availability (and cost) of MRI (Holliday et al., 2003) and remnant magnetisation (post-MRI) producing magnetic interference in MEG scans (Hutchinson et al., 2019; Kirschvink et al., 1992). For these reasons, the development of methods that avoid MRI is attractive.

The importance of avoiding MRI scanning is amplified further in wearable MEG. Nascent MEG systems comprising arrays of optically pumped magnetometers (OPMs; see Tierney et al. (2019), Brookes et al. (2022) and Schofield et al. (2022) for reviews) enable acquisition of MEG data from subjects who can move freely (Boto et al., 2018); even walk around a room (Holmes et al., 2023; Seymour et al., 2021). This has allowed data collection from an increased range of demographics, many of whom cannot tolerate conventional MEG systems, e.g. very young children (Boto et al., 2022; Corvilain et al., 2024; Feys et al., 2023, 2022; Hill et al., 2019; Rier et al., 2024; Vandewouw et al., 2024). While these participants could in principle be sedated for a structural MRI, this is unsuitable for scanning healthy participants in neuroscientific studies. There is therefore a risk that one of the major benefits of OPM-MEG – the ability to scan challenging cohorts – could be negated by the requirement for an anatomical MRI scan.

Template MRIs offer a potential solution. Recent years have seen the introduction of large databases of standard MRIs (Fillmore et al., 2015; Richards et al., 2016; Sanchez et al., 2012a, 2012b), which can ostensibly be used as approximations for individual anatomy. The use of template MRIs in source modelling using conventional MEG is common, with methods typically falling into four categories: 1) A single template is used for all participants in a study (Douw et al., 2018); 2) a template is selected via demographic matching (e.g., by age and/or sex, for example, López et al. 2014); 3) the ‘best-fitting’ MRI is selected from a database, by matching the head shape derived from the MRI to a 3D digitization of the participant’s head (Holliday et al., 2003; Seymour, 2018); 4) a pseudo-MRI is designed for each participant by warping a template MRI to the 3D digitisation of the scalp (Tadel et al., 2011). These methods have proved successful in conventional MEG. However, the ability to place sensors closer to the scalp means that OPMs offer MEG data with a higher signal strength (Boto et al., 2016; Hill et al., 2024; Iivanainen et al., 2017) and (in theory) better spatial resolution. In addition, OPMs measure neuromagnetic fields along multiple axes (Boto et al., 2022). Importantly, as data become more accurate, source analyses become more susceptible to inaccuracies in the forward model. This means that any approach to using template MRI scans with OPM-MEG data needs careful validation prior to deployment.

Here we describe a template warping method for use with OPM-MEG data. We use 3D structured-light scanning (Rocchini et al., 2001) – a method of 3D-image acquisition – to provide a high-resolution estimate of the subject’s head shape. Our method then takes age-matched template anatomical MRIs from an open-source database and warps them to the 3D head shape of the individual subject to generate a personalised “pseudo-MRI”. To test our approach, we acquire OPM-MEG data in a cohort of 20 healthy adult volunteers (all of whom have a real MRI scan – henceforth termed “individual-MRI”) and we undertake MEG analyses using equivalent pipelines but with the forward model informed either by the pseudo-MRI or the individual-MRI. We test a hypothesis that our pseudo-MRI approach will produce source space data that are highly correlated with the individual-MRI approach.

## Methods

### Pseudo-MRI generation

We used a 3D structured-light scanner (EinScan, H, Shining 3D, China) to estimate the size and shape of the participant’s head (Zetter et al., 2019). Briefly, the scanner projects a known pattern of light which is scattered by nearby objects and detected by a camera placed a known distance from the projector. As the structured light strikes a surface, any bumps or depressions cause a distortion of the reflected pattern recorded by the camera. The camera captures many such frames of data showing how different patterns of light are distorted, and by analysing these data it constructs point-clouds depicting the 3D surface of the object being scanned. Structured-light scanners have been commercialized and optimised for face and body scanning, meaning accurate 3D digitization can be acquired rapidly (in less than a minute) and comfortably. This technique is therefore suitable for use with participants who struggle to remain still and is considerably more practical (and cheaper) than MRI.

The method used to create a pseudo-MRI is summarized in Figure 1(a). First, a 3D structured-light scan is acquired while the participant is wearing an elastic cap to flatten their hair to the scalp surface; this enables an estimate of their head size and shape without hair getting in the way (Figure 1(b) – top left). Following this, an age-appropriate template anatomical T1-weighted MRI (i.e. an average of a number of age-matched MRI scans) is selected from a database (Richards et al., 2016) and a surface mesh depicting the scalp is extracted using FieldTrip (Oostenveld et al., 2011) (Figure 1b – top right). Meshes representing the outer head surface from both the structured-light scan and the template MRI are generated and a rigid body transformation is applied to co-register the two to the same space. Whilst in this space, the structured-light scan is cropped such that it covers similar areas of the head to the surface extracted from the MRI scan. The two meshes are then converted into binary 3D images, and “filled in” using a convex hull method (Figure 1b, middle row). This results in binary images that have a value of one inside the head and zero outside the head. Finally, FSL’s FLIRT(Jenkinson et al., 2002) is used to find the transformation matrix required to warp the binary images of the template MRI to the 3D structured-light image (parameters: X, Y, Z search from −90:90 degrees; correlation ratio cost function; 12 degrees of freedom; tri-linear interpolation). The derived transformation is then applied to the template MRI, generating a pseudo-MRI that has the same geometry as the outer head surface of the structured-light scan.

**Figure 1:**
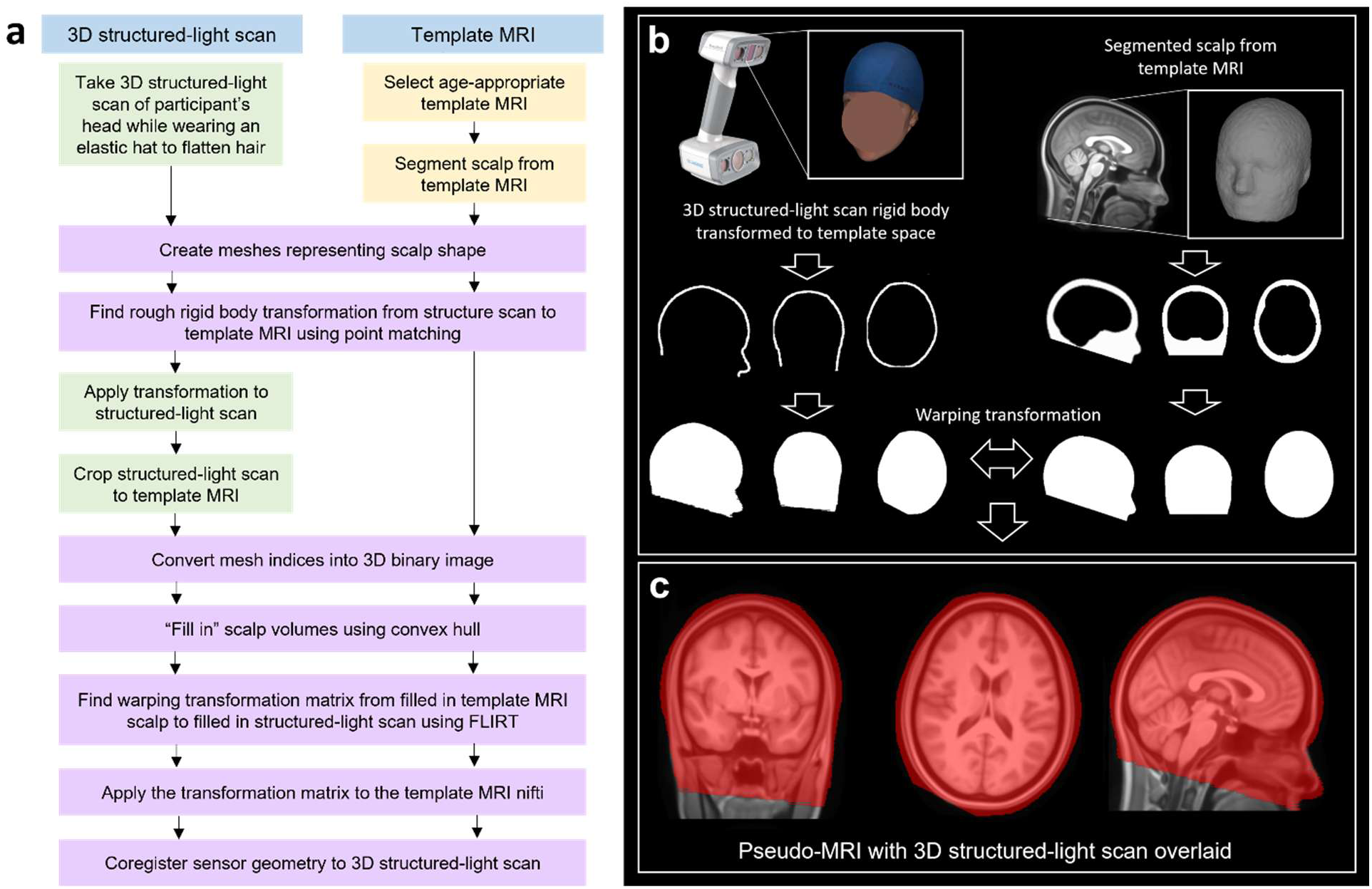
Generation of pseudo-MRI data: **a**) A flowchart describing the generation of pseudo-MRIs. **b**) Illustrations of real data demonstrating the method on a single participant. Top-left shows the structured-light scan and top-right shows the MRI. Both are used to create a binary image of the head, and the MRI is warped to fit the real subjects head shape from the structured-light scan. **c**) The final warped template anatomy (the pseudo-MRI) with the 3D structured-light scan overlaid in red.

A final result of this process is shown in Figure 1(c); here the underlying grey-scale image of the brain is the pseudo-MRI, and the red transparent overlay is the binary image based on the subjects’ real head shape. Note that all MRI templates were selected from the Neurodevelopmental MRI database (Richards et al., 2016).

### Data collection

#### MEG participants and paradigm

Twenty healthy adult participants took part in the study (age range 23 – 34; mean age 27; 10 identified as male, 10 as female). All participants provided written informed consent prior to data acquisition and the study was approved by the University of Nottingham’s Faculty of Medicine and Health Sciences Research Ethics Committee. These data – which were collected as part of a neurodevelopmental study – have been previously published (Rier et al. 2024).

The task is outlined in Figure 2 and involved sensory stimulation. A single stimulator (Metec, Germany) comprised 8 independently controlled “pins” which could be raised or lowered (using a piezo-electric crystal) to tap the participant’s finger. A single trial comprised 0.5 s of stimulation (during which the finger was tapped 3 times using all 8 pins) followed by 3 s rest. We used two separate simulators to deliver stimulation to either the index or little fingers; the finger stimulated was alternated between trials. There was a total of 41 trials for each finger (82 trials in total) and the experiment lasted 287 s. Throughout the experiment, subjects were seated comfortably on a patient support and watched a television program of their choice (presented via back projection onto a screen in the MSR located about 1 m in front of the subject). Subjects were free to move their head during the experiment although they were not encouraged to do so.

**Figure 2:**
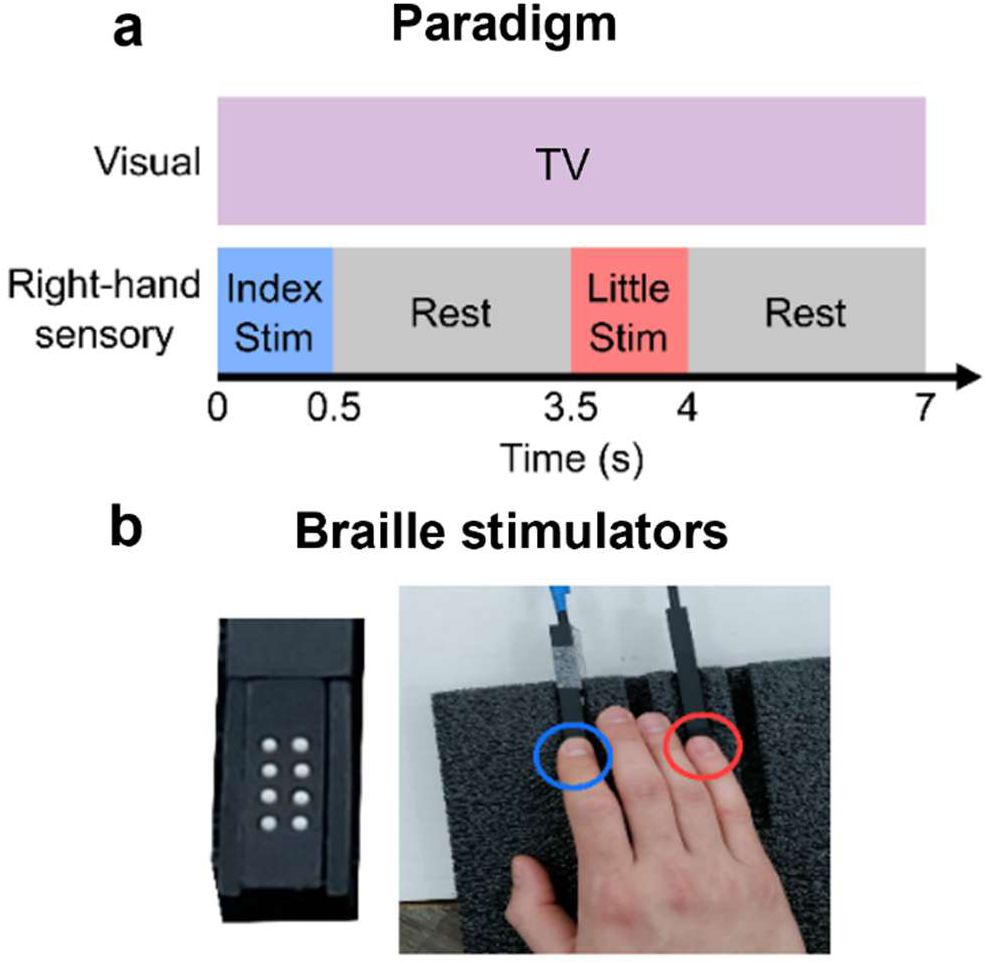
Sensory paradigm: **a**) A timeline of the sensory paradigm. **b**) Images showing the braille stimulators (left) and hand placement over the two stimulators on the index and little finger of the right hand (right). This figure is adapted from Rier, et al. (2024).

#### Imaging system

The OPM-MEG system housed up to 64 triaxial sensors (3^rd^ generation OPMs, QuSpin Inc., Boulder, CO, USA) each capable of measuring magnetic field in three orientations. The array could therefore acquire data via up to 192 independent channels. (Note that, due to the experimental nature of the system, not all sensors were available for all scans and so channel count varied between subjects.) Sensors were uniformly distributed across rigid 3D-printed helmets that came in multiple sizes to accommodate changes in head size (Cerca Magnetics Limited, Nottingham, UK). All sensors were synchronized, and their analogue outputs sampled at 1,200 Hz using a National Instruments (NI, Texas, US) data acquisition system interfaced with LabView (NI). The system was housed in an OPM-optimised magnetically shielded room (MSR) comprising 4 layers of mu-metal, a single layer of copper and equipped with degaussing coils (Magnetic Shields Limited, Kent, UK); the inner walls of the room were degaussed prior to every scan to reduce remnant magnetisation (Altarev et al., 2014). The environmental static magnetic field was further suppressed using a field nulling technique in which the known background field in the MSR was measured, and then reduced by applying an equal and opposite field delivered using a set of electromagnetic coils (Holmes et al., 2018; Rea et al., 2022, 2021; Rhodes et al., 2023).

Immediately following MEG data acquisition, two 3D digitization’s of the participants’ head, with and without the OPM helmet, were generated (using 3D structured-light scanning). These were used for both pseudo-MRI generation (see above) and coregistration of the OPM sensor locations and orientations to brain anatomy.

For all subjects, a volumetric anatomical MRI scan was acquired using a Phillips Ingenia 3T MRI system (running an MPRAGE sequence with 1 mm isotropic resolution and *T*_1_ contrast).

### Data analysis

We processed all 20 OPM-MEG datasets twice, once using the individual-MRI and a second time using the pseudo-MRI. The two pipelines were identical (unless otherwise stated) and independent (i.e. the individual-MRI wasn’t used in the pseudo-MRI pipeline, and vice versa).

#### Coregistration

For *coregistration to the pseudo-MRI*: we aligned a 3D mesh of the OPM-MEG helmet (from which the sensor locations/orientations are known) to the structured-light scan of the participant wearing the helmet; we then aligned the facial features from the same scan, to the structured-light scan of the participant without the helmet on; this second scan (with no helmet) was already co-registered to the pseudo-MRI, and this therefore allowed complete coregistration of the sensor array geometry to the pseudo-MRI.

For *coregistration to the individual-MRI*: we took the 3D image of the participant without the helmet and aligned the facial features to those extracted from their individual MRI (using manual feature selection and the iterative closest point algorithm in MeshLab (Cignoni et al., 2008)); we then combined this with the transform obtained above, from the 3D mesh of the OPM-MEG helmet to the 3D image of the participant without the helmet, and this enabled a complete coregistration of the sensor array geometry to the individual-MRI.

#### Quantitative comparison of brain anatomy

Initially we aimed to quantify the difference in brain anatomy between individual-MRI and pseudo-MRI. To this end, we employed the automated anatomical labelling (AAL) atlas (Gong et al., 2009; Hillebrand et al., 2016; Tzourio-Mazoyer et al., 2002). For both the individual and pseudo MRI’s: 1) the brain was extracted from the MRI using FieldTrip (Oostenveld et al., 2011); 2) the MNI standard brain was co-registered to the individual anatomy; 3) the same transform was applied to the AAL region map. (These steps were completed using FLIRT in FSL (Jenkinson et al., 2002; Jenkinson and Smith, 2001)). 4) The medoid of each AAL region was then found. This resulted in two sets of AAL coordinates: one for the individual-MRI and one for the pseudo-MRI. Both the individual-MRI and pseudo-MRI were in the same coordinate frame (i.e. they were defined relative to the MEG sensor locations), and this processes therefore allowed us to calculate the Euclidean distances between AAL centroids derived using the two methods. This provides a quantification of the differences in brain shape between the real anatomy and the estimated (pseudo) brain anatomy.

#### MEG Data pre-processing

For MEG data, we used a pre-processing pipeline described previously (Rier et al., 2023). Broken or excessively noisy channels were identified by visual inspection of channel power spectra and removed. Notch filters at the powerline frequency (50 Hz) and 2 harmonics, and a 1 – 150 Hz non-causal 4^th^ order Butterworth bandpass filter, were applied. Bad trials were defined as those with variance greater than 3 standard deviations from the mean trial variance, and automatically removed. A visual inspection was also carried out and any remaining trials with excess artifacts removed. Eye blink and cardiac artefacts were removed using ICA (implemented in FieldTrip (Oostenveld et al., 2011)) and homogeneous field correction (HFC) was applied to reduce interference that manifests as a homogeneous field across the helmet (Tierney et al., 2021). On average (across participants) we had 159 ± 9 available channels, and we removed 4 ± 2 trials. (Note the large variation in channel count was primarily a result of different sensor counts being available for different subjects, rather than rejection of bad channels.)

#### Measuring beta modulation

The modulation of beta band amplitude during index and little finger stimulation was analysed separately for the two trial types. Data were filtered to the beta band (13 – 13 Hz) using a non-causal 4^th^ order Butterworth bandpass filter. Source localisation was performed using a linearly constrained minimum variance (LCMV) beamformer (Robinson and Vrba, 1998). We generated beamformer images of the spatial signature of beta modulation using two approaches:

- A whole brain analysis, in which we divided the brain into 4 mm isotropic voxels.
- A “high-resolution” analysis where we masked the sensorimotor cortices (by taking the left pre- and post-central gyri from the AAL atlas, dilating these volumes using a 5 mm sphere, and then masking those volumes for all subjects). Brain regions inside the mask were divided into 1 mm cubic voxels.

In both cases, forward solutions were computed using a single shell model (Nolte, 2003). Covariance matrices were generated using beta band filtered data and separately for index and little finger trials (excluding bad trials). All covariance matrices were regularized using the Tikhonov method with a regularization parameter equal to 5 % of the maximum eigenvalue of the unregularized matrix (Brookes et al., 2008). The optimised source orientation for each voxel was taken as that with the largest projected power (Sekihara et al., 2004). Beamformer weights were generated for each voxel. Pseudo-T statistical images, contrasting source power in active (0.3 – 0.8 s relative to movement onset) and control (2.5 – 3 s) windows, were generated separately for the index finger and little finger trials. This analysis was carried out independently using the individual MRI and pseudo-MRI, resulting in 8 images per subject for the following conditions: index and little finger; low and high resolution, and individual- and pseudo-MRI.

Having identified the location in the brain of peak stimulus-induced beta modulation (based on the high-resolution maps), a broadband (1 – 150 Hz) estimate of electrophysiological activity at this peak location (termed a virtual electrode measurement) was calculated. This was computed using a beamformer (with covariance based on broadband data, but otherwise implemented as above). The time-frequency content of this signal was analysed using a Hilbert transform. Briefly, the data were filtered into multiple overlapping frequency bands; within each band, the analytic signal was derived based on a Hilbert transform, and its absolute value computed giving the amplitude envelope of beta oscillations. These envelopes were averaged across trials. A baseline (defined in the 2.5 s to 3 s window) was subtracted, and the resulting data were normalised by the same baseline to generate a time-frequency decomposition showing relative change in oscillatory amplitude from baseline. This process resulted in 4 time-frequency plots per subject (index/little finger trials, and individual/pseudo-MRI).

#### Measuring Connectivity

Whole-brain connectivity was quantified using amplitude envelope correlation (AEC) (Brookes et al., 2011; Liu et al., 2010). Beta band filtered regional signals from the 78 AAL regions were calculated using a beamformer (implemented as above with beta band covariance). A time window, 0.1 s - 3.4 s relative to trial onset was selected to exclude edge effects and all trials were concatenated. Source leakage was corrected using pairwise orthogonalisation (Brookes et al., 2012). The absolute value of the Hilbert transform was computed to generate the amplitude envelope from the oscillatory signal. Envelopes were downsampled to 120 Hz and the Pearson correlation computed between all possible AAL region pairs, generating a 78 x 78 matrix for each participant describing the beta band connectivity. The “global connectivity” in the beta band for each participant was calculated by summing all off-diagonal elements of the connectivity matrix. The connectomes from the pseudo and individual MRIs were compared across individuals, the group average, and regional averages using Pearson correlation.

#### Removing the effects of coregistration error

The above analyses directly compared individual- and pseudo-MRI approaches. However, they also conflate multiple sources of error:

- **Volume conductor error:** Error in the forward field model due to having a different volume conductor model (the real brain/head shape versus an estimated brain)
- **Coregistration error:** Differences in the way coregistration is achieved for the individual and pseudo-MRI approaches will lead to two slightly different co-registrations.
- **AAL region definition:** Differences in brain shape will necessarily lead to the AAL centroids being in slightly different positions for the individual and pseudo-MRI approaches (though this only affects connectivity analysis).

In a situation where pseudo-MRIs are used (and no real MRI scan is available) conflation of these errors is unavoidable. However, as real MRIs were available in this study, some of these errors can be disentangled. To this end, we rigidly aligned the individual MRI brain to the pseudo-MRI using FLIRT (Jenkinson et al., 2002), meaning we could apply the same coregistration to both analyses. We then repeated the above analyses (of beta band modulation and connectivity) using MRIs in the same space and using the same coregistration, thus eliminating the errors due to coregistration. Individual results with and without coregistration error were plotted in histograms and Gaussians fitted to the binned data.

## Results

### AAL comparison

We compared the medoids of the 78 cortical regions from the AAL atlas, derived using the individual and pseudo-MRIs. Figure 3(a) shows a single representative participant, in whom the median distance between regions was 2.54 mm. Across all 20 participants, at the individual subject level the medoids were separated by an average of 2.46 ± 0.42 mm (note here, the median Euclidean distances across locations was computed separately for each subject, and the median and median absolute deviation across subjects presented). At the group level, the average distance between medoids (calculated by first averaging locations across subjects by taking the mean, and then computing the median location discrepancy across regions) was 1.1 mm.

**Figure 3.**
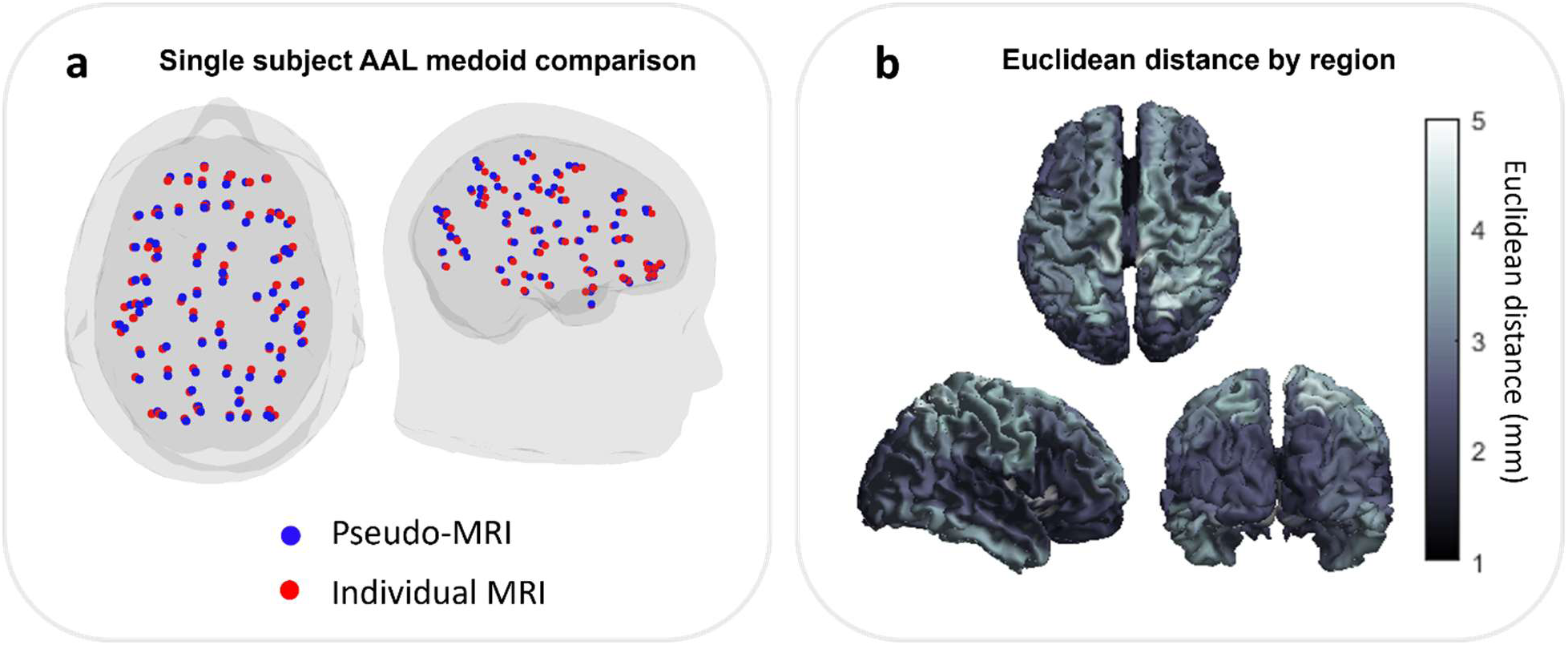
**a**) an example participant with the location of the medoids of 78 cortical regions as dictated by the AAL atlas, derived from the pseudo-MRI (blue) and individual MRI (red). **b**) distance between medoid location by cortical region, averaged across the 20 participants.

Figure 3(b) shows the median distances calculated (at the individual level) for all 78 AAL regions independently. Dorsal regions tend to show a larger Euclidean distance than frontal regions. This will be addressed in the discussion.

### Beta modulation

Figures 4(a) and 4(b) show beta band modulation for an example participant during the sensory task; panel (a) shows index finger stimulation and panel (b) shows little finger stimulation. In both cases the upper panel shows the pseudo-T statistical images contrasting stimulation and rest windows, overlaid on a standard brain. The left-hand side shows results from the individual MRI approach and the right-hand side shows the pseudo-MRI approach. The beta modulation (shown by more negative pseudo-T values) localises to the left sensorimotor cortex using both models, as would be expected. For this individual, the peak locations for index finger stimulation, derived using our two methods, were separated by 14.99 mm; the separation of peak locations for little finger stimulation was 11.84 mm. The median separation across all 20 participants was 11.46 ± 3.39 mm for index stimulation and 11.60 ± 3.72 mm for little finger stimulation (median ± median absolute deviation). The Pearson correlation between the (vectorised) pseudo-T statistical images for this participant was 0.87 for index finger stimulation and 0.85 for little finger stimulation. The median correlation between images across the 20 participants was 0.76 ± 0.07 for index finger stimulation and 0.74 ± 0.06 for little finger stimulation. These relatively high correlations demonstrate that similar results can be generated with or without an MRI scan.

**Figure 4:**
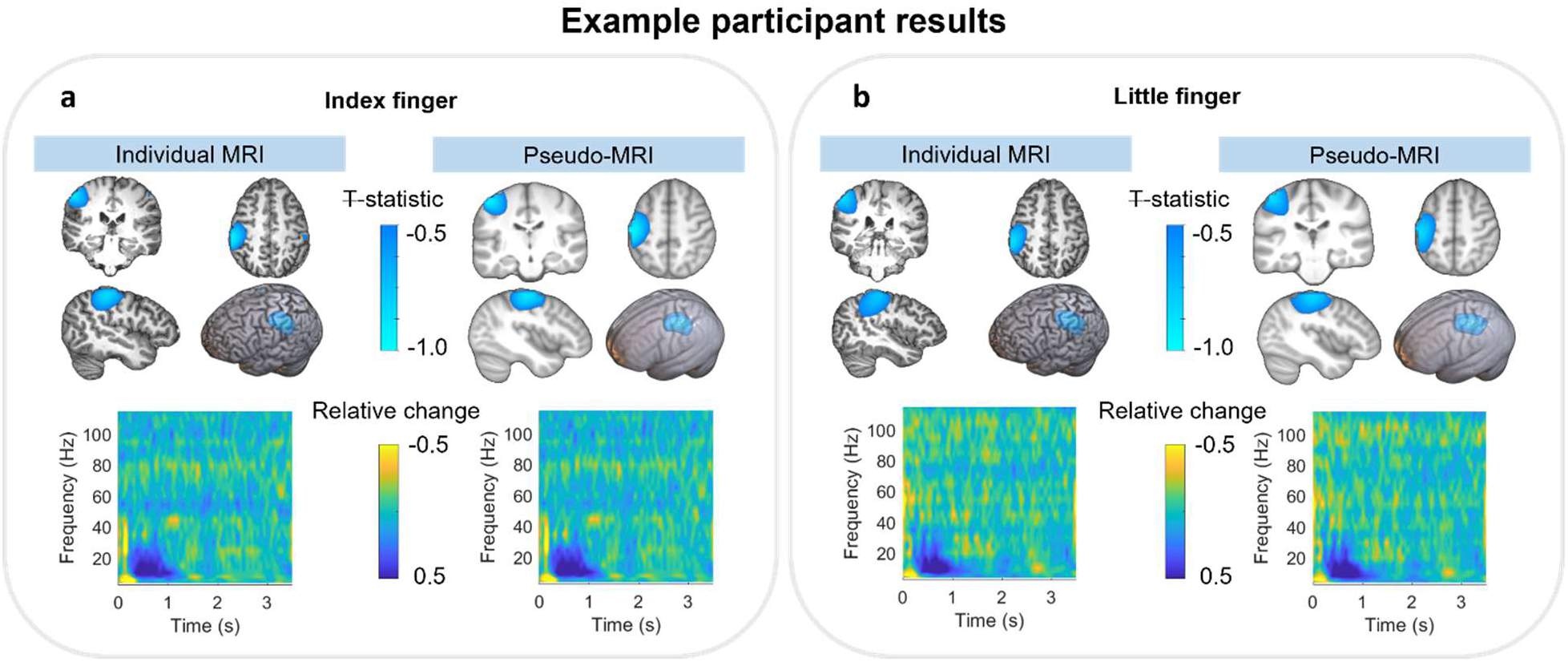
Beta modulation for an individual subject. (a) and (b) show results for the index and little finger stimulation respectively. In both cases, the upper panel shows pseudo-T statistical maps depicting the spatial signature of task induced beta modulation across the brain; these images are derived from either the individual MRI (left) or the pseudo-MRI (right). Maps are overlaid on the individual and pseudo-MRI extracted brains. The time-frequency spectrograms below show the evolution of spectral power throughout the average task trial.

The lower panels of Figures 4(a) and 4(b) show time frequency spectra from the location of maximum beta modulation; yellow represents an increase in oscillatory power and blue represents a decrease. As expected, when using either the individual or pseudo-MRI approach we see a drop in beta band power in the 0-1 s window (i.e. during finger movement). The Pearson correlations between TFS’s, derived using the two methods for this example subject were 0.97 for index finger stimulation and 0.94 for little finger stimulation. The median correlations across 20 participants were 0.97 ± 0.01 for index finger stimulation and 0.95 ± 0.03 for little finger stimulation.

Figure 5 shows the group averaged results across 20 participants. The Euclidean distance between peak locations from the group average pseudo-T statistical maps was 2.75 mm and 3.73 mm for index and little finger, respectively. The Pearson correlation coefficients between group average individual and pseudo-MRI derived pseudo-T statistical maps are 0.95 and 0.92 for index finger and little finger stimulation, respectively. Group average time-frequency spectrograms (TFS) from virtual electrodes at the location of the largest beta modulation are shown below the pseudo-T statistical maps. The Pearson correlations between the corresponding TFS are 0.99 and 0.98.

**Figure 5:**
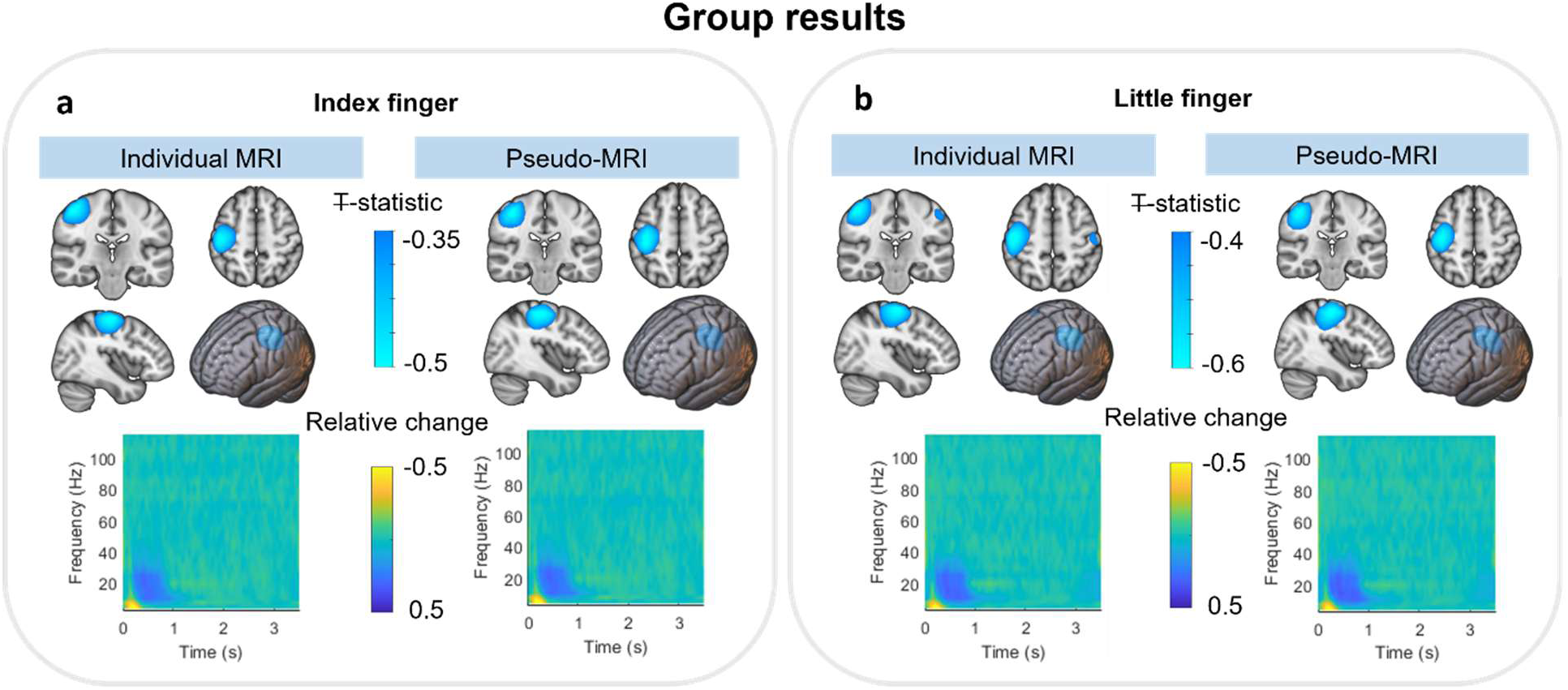
Group averaged beta results. (a) and (b) show results for the index and little finger stimulation respectively. In both cases, the upper panel shows pseudo-T statistical maps depicting the spatial signature of task induced beta modulation across the brain; these images are derived from either the individual MRI (left) or the pseudo-MRI (right). Maps are overlaid on the individual and pseudo-MRI extracted brains. The time-frequency spectrograms below show the evolution of spectral power throughout the average task trial.

### Connectivity

Figure 6 shows the results from the group level connectivity analysis. For these analyses, both index and little finger trials were concatenated, as in Rier et al. (2024). Panel (a) shows the group average connectivity matrices from the individual MRI analysis (left) and pseudo-MRI analysis (right). Panel (b) shows the corresponding glass brain plots displaying the top 5% of connections (i.e.,5% of all connections with the highest connectivity values, therefore showing the 150 strongest connections) for the individual (left) and pseudo-MRIs (right). The blue circles on the glass brain are scaled by connectivity strength (i.e. how connected each node is to all other nodes; in other words, the sum of each row of the matrix). Both plots show a bilateral sensorimotor network, as expected during the sensory stimulation.

**Figure 6.**
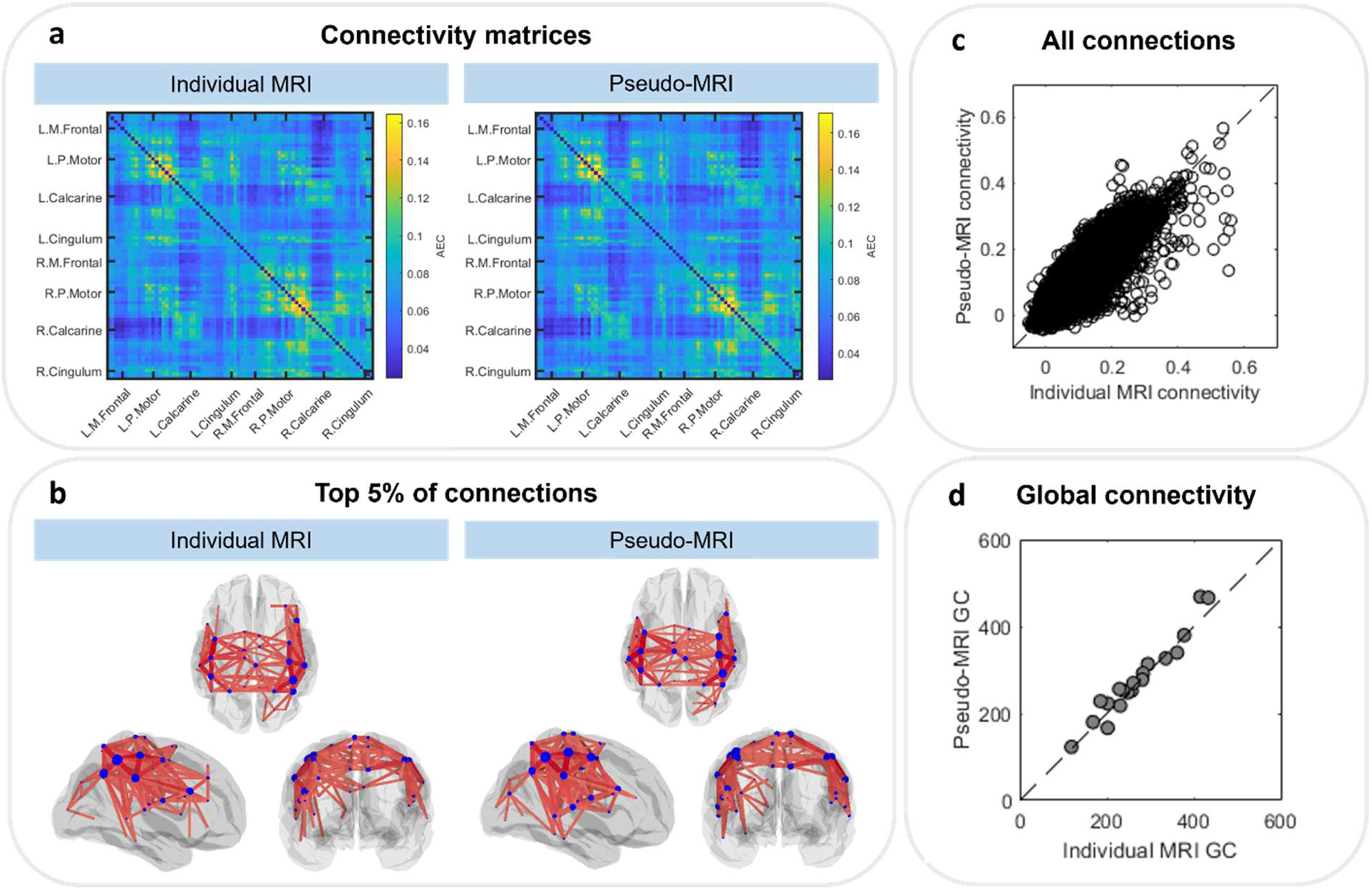
**a**) group average connectivity matrices derived from analysis with the individual MRI anatomy (left) and pseudo-MRI (right). **b**) the top 5% of connections overlaid on glass brains derived from analysis with the individual MRI anatomy (left) and pseudo-MRI (right). **c**) The relationship between all connections from all participants. **d**) the relation between global connectivity (sum of off-diagonal elements of the connectivity matrices) for each participant from the individual MRI and pseudo-MRI.

The relationship between all connections for all participants is explored in panel (c), which shows the values of the elements in the two matrices in panel (a) plotted against each other. The result is a Pearson correlation coefficient of 0.87. We also computed global connectivity (i.e., the sum of connectivity across all region pairs) for each individual subject and Panel (d) shows these values computed using the individual-MRI plotted against the same values derived using the pseudo-MRI; the Pearson correlation is 0.98. Finally, we measured the Pearson correlation between connectivity matrices at the individual level, which had a median value across 20 subjects of 0.82 ± 0.03.

### Removing coregistration error

Figure 7(a) shows the distribution of median Euclidean distance between AAL regions identified from the individual and pseudo-MRI approaches. The distribution is computed across participant’s and a Gaussian is fitted to the median value (of 2.46 mm). Figures 7(b) and 7(c) show histograms describing the individual participants beta modulation for index and little finger stimulation, respectively, again with Gaussian fits overlaid. In both cases the upper plot shows the spatial discrepancy between peak locations using individual and pseudo-MRI’s; the middle plot shows the correlation between the pseudo-T-statistical images and the lower plot shows the correlation between TFS’s from the peak location. In all cases, orange represents the results with independent coregistration using the two separate MRI’s (i.e. with coregistration error), and purple shows the equivalent data with coregistration error eliminated. Note that coregistration has a large effect on the pseudo-T-statistical images, yet a relatively small effect on the TFSs, which were highly correlated despite the presence of coregistration errors.

**Figure 7.**
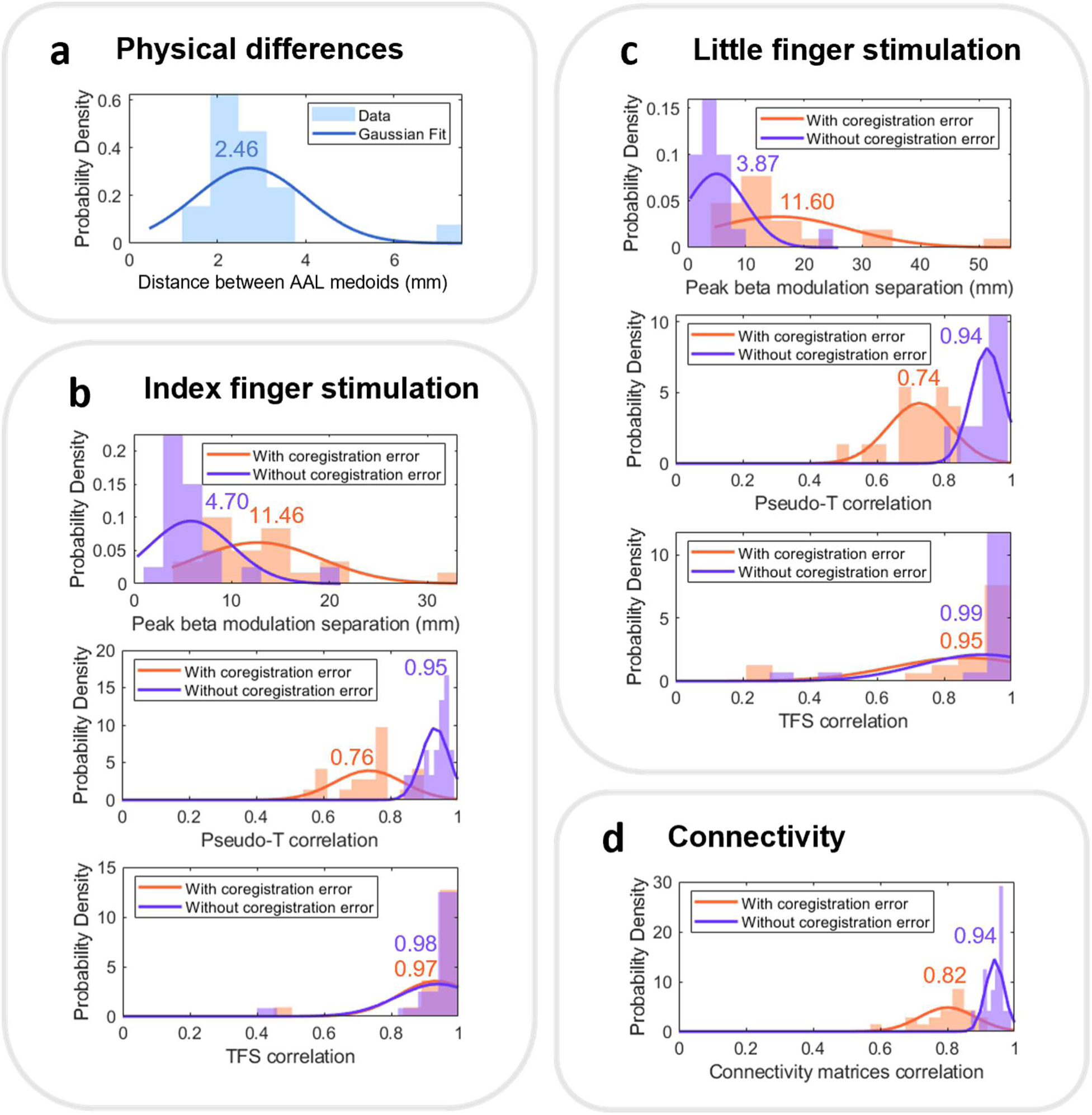
Histograms summarising the results across the 20 participants. **a)** shows the Euclidean distance between AAL medoids from individual and pseudo-MRI anatomy. **b-c)** show the results of the index and little finger stimulation, respectively. **d)** describes the connectivity results. For b-d, orange describes the results from analyses using separate co-registration for individual and pseudo-MRI (compounding co-registration errors) and purple shows the repeated analyses following transformation of the individual MRI to the pseudo-MRI, using the same coregistration. Inset values show the median value.

Finally, Figure 7(d) shows the correlation between beta band connectivity matrices with and without coregistration error. We find that eliminating coregistration error increases the correlation between correlation matrices.

## Discussion

Wearable OPM-MEG opens new avenues for neuroscientific research, by enabling researchers to study participant groups that find a conventional neuroimaging environment challenging (e.g. where participants must remain still for extended periods). However, the value of this is limited if MRI scans (which also require participants to remain still) are required. Here, we have shown that a method of warping template MRIs to 3D structured-light scans of the head to generate pseudo-MRIs produces similar MEG results to those generated if individual MRI scans are used. This suggests that, particularly for large neuroscientific group studies, our pseudo-MRI approach might be preferable than acquiring individual MRIs for every participant.

We tested our method using data collected on 20 adult participants, first examining the Euclidean distances between cortical regions identified using individual anatomy and pseudo-MRI. This is essentially a quantification of how different the locations of anatomical landmarks are, when identified using the real and the pseudo-MRI. On average we found good agreement, with a separation of 2.46 ± 0.42 mm at the individual subject level). The parietal cortex had the greatest discrepancy, and this may be due to the 3D digitisation being less representative of the true head shape at the top of the head, due to hair, and the elastic swim cap. This could be mitigated by using tighter-fit elastic caps or by accounting for hair thickness when generating the pseudo-MRI.

We examined the sensitivity of MEG analyses to the pseudo-MRI by analysing data recorded during a somatosensory task. Specifically, we used two equivalent analysis pipelines, with forward models generated by individual and pseudo-MRI, and in both cases, we found beta modulation during right-hand sensory stimulation localised to the left sensorimotor cortex. As in Rier et al. (2024), we did not find separation between index and little finger representations; this is likely because the experiment was not optimised for this. The distance between voxels showing the highest beta modulation was, on average, around 11 mm at the individual subject level; this compares well to the study by Douw et al. (2018) which used template MRIs and conventional MEG data. Related, the correlation between pseudo-T statistical maps was ~0.75, suggesting that the spatial patterns of beta activity are similar despite the difference in forward model. At the group level the discrepancy between the two approaches was reduced; the median peak voxel locations were separated by just <3 mm and the correlation between images was ~0.95, implying that the differences between real- and pseudo-MRIs at the individual level are likely random and so average out across a group.

We also explored the impact of pseudo-MRI anatomy on functional connectivity. Specifically, we derived beta-band connectomes between 78 cortical regions and found high consistency at the group level with a correlation of 0.87 between connectivity matrices derived using the two methods. Individual global connectivity was also consistent between analyses, and the individual connectome matrices showed a correlation of >0.8.

The above results were all derived using completely independent pipelines (i.e. the pseudo-MRI analyses never used the real MRI and vice versa). However, this conflates 3 sources of error; 1) **Coregistration error**: i.e. coregistration of sensor geometry to brain anatomy is performed independently using two MRI scans and so the sensor locations and orientations will differ between the two analyses. 2) **Forward field error**: the brains/heads are diffe4rent shapes/sizes and this will lead to a difference in the volume conductor model used to derive the forward field. 3) **AAL region error**: because the brains are different shapes/sizes, the AAL locations are in slightly different positions (which will affect connectivity analysis). In this study we were able to coregister the two MRI scans, and in doing so remove the effect of coregistration error; importantly this would be impossible in a real study (as no individual MRI’s would exist), but for our purposes showed that the likely largest source of discrepancies between our two approaches was coregistration error. This is an important finding and something that should be the topic of future work.

There are a few notable limitations of the method described here. Firstly, the template MRIs, while approximately age-matched (using 5-year windows), were not sex-matched. As there may be structural differences in the brains of males and females (Giedd et al., 2012) beyond just head size, accounting for this might improve the overall results. Second, how this method applies across diverse participant groups should be considered, as it requires hair styles that can flatten to get an accurate head shape estimate. Third, we used a single shell forward model, as this has been suggested to be comparable to more complex BEM models (Stenroos et al., 2014). However, this model may be less susceptible to inaccuracies in anatomy than, for example, a three-shell BEM. This should be investigated in future work.

## Conclusion

We have demonstrated a method for warping template anatomical MRIs to participants’ head shapes derived from 3D structured-light scanning. We have shown evidence that pseudo-MRIs perform comparably to individual MRI anatomy in a range of typical MEG analyses. This method will allow source reconstruction of MEG data from participants who typically struggle with the MRI scanning environment. This is of particular importance with for wearable MEG using OPM sensors.

## Data and code availability

All data used to produce the results presented here are published in (Rier et al., 2024) and made available on Zenodo (https://zenodo.org/records/11126593). All code was made available on GitHub (https://github.com/nsrhodes/template_warping).

## Author contributions

**NR**: Conceptualization, Methodology, Data curation, Formal analysis, Software, Writing – original draft; **LR**: Conceptualization, Methodology, Data curation, Software, Writing – reviewing and editing; **EB**: Methodology, Supervision, Writing – reviewing and editing; **RMH**: Methodology, Software, Writing – reviewing and editing; **MJB**: Conceptualization, Funding acquisition, Supervision, Writing – reviewing and editing.

## Funding

This work was supported by an Engineering and Physical Sciences Research Council (EPSRC) Healthcare Impact Partnership Grant (EP/V047264/1) and an Innovate UK germinator award (Grant number 1003346). We also acknowledge support from the UK Quantum Technology Hub in Sensing and Timing, funded by EPSRC (EP/T001046/1).

## Declaration of competing interests

MJB is a co-founder with equity and the chairman of Cerca Magnetics Ltd., who are commercialising OPM-MEG technology. E.B. is the chief technology officer of Cerca Magnetics Ltd, with equity. R.H. is a scientific advisor for Cerca Magnetics Ltd, with equity. L.R. is a software consultant for Cerca Magnetics Ltd.

